# Spatial single-cell interactome and niche-specific molecular signatures in alcohol-related liver disease

**DOI:** 10.1101/2025.03.15.643421

**Authors:** Oleksandr Petrenko, Thomas Sorz-Nechay, Martin Bilban, Sophia Derdak, Lukas Van Melkebeke, Wenxin Huang, Jef Verbeek, Schalk van der Merwe, Benedikt S Hofer, Benedikt Simbrunner, Thomas Reiberger, Tim Hendrikx

**Author notes:** Correspondence should be addressed to T.H.

## Abstract

Alcohol-related liver disease (ALD) remains a major global health burden with limited therapeutic options due to an incomplete understanding of its underlying molecular mechanisms and cellular crosstalk. Here, we applied ultra-high resolution (on 2 µm spots) spatial transcriptomics to a cirrhotic liver tissue obtained from an end-stage ALD patient, analyzing >265,000 spatially resolved cells with further validation on single-cell and single-nuclei datasets from patients with ALD cirrhosis. Our analysis delineated distinct cellular sub-populations and molecular landscapes across fibrotic, vascular, and parenchymal niches of ALD cirrhosis. We identified robust zonation of hepatocytes, hepatic stellate cells, and diverse immune subpopulations, including enrichment of pro-inflammatory T cells and dendritic cells in the fibrotic niche and *MARCO*+ tissue-resident macrophages localizing mostly in parenchymal areas. Analysis of spatial metrics assigned expression of *WNT4, RCAN3, PPIAL4G, PLA2G5*, and *SLC6A9* to the fibrotic environment in ALD. Differential expression and ligand–receptor interactome analyses revealed niche-specific signaling, with marked *CCL19*–*CCR7* activity in fibrotic regions and *DLL4*–*NOTCH3* crosstalk in vascular compartments. Notably, *WNT4*+ fibroblasts emerged as key mediators of extracellular matrix remodeling and chemoattraction, particularly via CCL19-mediated signaling towards CD8^+^ T cells, which was validated on single-cell resolution within the ALD cirrhotic liver in external datasets. These spatial and single-cell findings highlight novel potential therapeutic targets for patients with ALD cirrhosis.

## Introduction

More than 3% of the global population is affected by alcohol-related liver disease (ALD), and up to 25% of individuals with harmful alcohol consumption will develop liver cirrhosis, associated with life-threatening complications and a 5-year mortality rate of ∼60%^1,2^. The histopathological features of ALD are characterized by a centrilobular injury pattern featuring more often macrovesicular steatosis, hepatocellular ballooning, and lobular inflammation, with early fibrotic changes manifesting as perivenular and perisinusoidal fibrosis^3,4^. With continued excessive alcohol intake, these lesions can progress to bridging fibrosis and, ultimately, cirrhosis, where the architecture of the liver is severely disrupted^2,5^. While abstinence facilitates partial regression of ALD histopathology and fibrosis in earlier stages, advanced cirrhosis is associated with high morbidity and an increased risk of decompensation^6–8^. Ultimately, there is no effective pharmacological treatment available to halt the progression of ALD or to induce fibrosis regression. Given the enormous societal and economic burden and the shortage of donor livers, with liver transplantations only possible at given medical centers, a better understanding of the underlying molecular pathology is necessary to provide novel therapeutic options to improve the outcome of ALD patients.

In the recent past, efforts were undertaken to supplement the histopathological understanding of ALD with massively parallel transcriptional profiling in order to fill gaps in ALD biology and identify prospective molecular targets. The first such study on a single-cell resolution, published by P. Ramachandran et al., included, among other etiologies, 2 ALD cirrhosis datasets generated from explanted livers and described *TREM2*^+^*CD9*^+^ macrophage population associated with fibrotic niches during liver disease^9^. Similarly, a study by C. Gao et al. presented five single-cell datasets obtained from patients with ALD biopsies, focusing on T-cell diversity and describing *GZMK*^+^*CD4*^+^ T-cell infiltrate as a characteristic of ALD^10^. Recent developments in spatial biology, however, allowed profiling of cellular transcriptomes on a tissue slide, focusing not only on “what” significant processes occur in liver fibrosis but also “where” they are situated in the tissue microenvironment. As such, the presence and localization of earlier discovered *TREM2*^+^ macrophages at fibrotic areas in the steatotic liver was shown by us^11^, besides experimental evidence describing their functional role in scar remodeling. Moreover, this led to the identification of systemic soluble TREM2 levels as an indicator for the degree of liver disease, altogether demonstrating the significant usability of spatially resolved transcriptomic profiles at the single-cell level. Such valuable readouts are yet to be available in the field of chronic liver disease and, specifically, in ALD.

Here, we employed ultra-high resolution (2µm spots) spatial transcriptomics to comprehensively profile the cellular landscape of a cirrhotic liver from a patient with endstage ALD. We report cellular populations, their functional sub-types, transcriptomic markers, and cellular interactions linked to ALD cirrhosis, as well as the validation results of these findings on single-cell resolution in an external dataset.

## Results

### Spatial transcriptome analysis at ultra-high resolution identified various prevalence and zonation of immune infiltration

The patient (male, 54 years old) enrolled in this study was diagnosed with end-stage alcohol-related liver disease and was scheduled for liver transplantation at the Vienna General Hospital, Austria. Previous histopathological findings included liver cirrhosis (Figure 1A), mixed steatosis, and pronounced immune infiltration. MELD-Na score before liver transplantation was 10 points, Child-Pugh score was 10 points, with laboratory findings: ALT = 787 units/L, AST = 1132 units/L, bilirubin = 1.32 mg/dl, INR = 1.4, platelets = 68 G/L. Following the collection of a sample of the cirrhotic liver during transplant surgery, the obtained sample was stained with picrosirius red, and digital quantification of the collagen fiber showed a 33.4% collagen proportionate area. There was co-localization of the fibrosis area with activated hepatic stellate cells (HSC) as shown by alpha-smooth muscle actin staining (positive area = 28.2%, Supplementary Figure 1A, Supplementary Files 1, 2).

**Figure 1.**
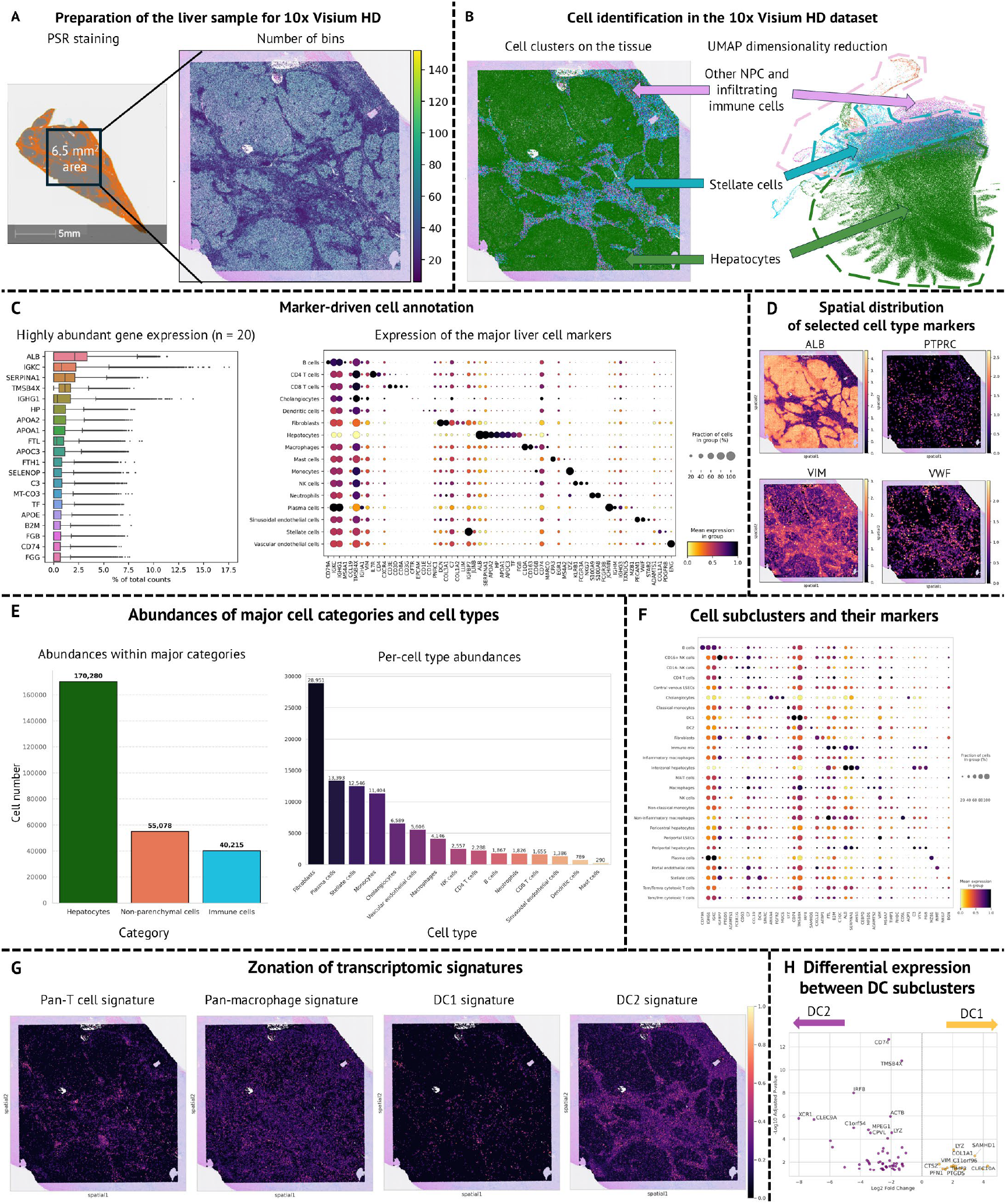
Spatial transcriptomics in alcohol-related liver disease allowed the identification of major cell types and their sub-populations. **(A)** The sequencing area was selected based on PSR staining to capture fibrotic, vascular, and parenchymal areas of the liver. **(B)** Clustering of the cells based on their transcriptomes identified distinct cell populations, separated both on a dimensionality reduction plot and when transferring labels to the tissue slide. **(C)** After filtering, classical cell markers such as ALB, IGHG1, and CD74 were among the most abundantly expressed genes. Furthermore, marker-driven annotation identified all major liver cell populations. **(D)** Expression of the selected cell markers display zonation of hepatocytes (ALB), immune cells (PTPRC), mesenchymal-lineage cells (VIM) and endothelium (VWF). **(E)** The quantification of the annotated cells showed that hepatocytes were the predominant cell population, followed by comparable counts of non-parenchymal and immune cells. Cell-type-specific quantification showed that fibroblasts/stellate cells, B cells and T cells, and monocytes were the most abundant within the dataset. **(F)** Application of the CellTypist algorithm with a model trained on human liver cells allowed for increased resolution, identifying cell sub-populations such as two dendritic cell clusters. **(G)** Display of selected transcriptional signatures (gene modules) with their expression strength in regard to tissue zones. **(H)** Volcano plot displaying differentially expressed genes between two DC subclusters. Only genes meeting significance thresholds after differential expression with t-test (i.e., log2(fold change) > |1|, Padjusted < 0.05) are displayed, and 20 genes with the highest rank are labeled. Abbreviations: PSR = picrosirius red. NPC = non-parenchymal cells. DC = dendritic

High definition (HD) spatial transcriptomics resulted in recovery of 10.836.960 2µm spots, further combined in 265.573 cells post-quality control. Overall, fibrotic areas contained less accessible spots than non-scarred parenchyma (Figure 1A). An unbiased graph-based clustering on whole transcriptomes confirmed the separation of cells in distinct populations expressing markers of hepatocytes, stellate cells, other non-parenchymal (NPC), and immune cells (Figure 1B, Supplementary Data 1). As expected, cells identified as HSC were predominantly found in fibrotic areas, while most of the parenchyma were expectedly identified as hepatocytes (Figure 1B). Various cellular markers were identified among the highest-expressed genes in the dataset: albumin (*ALB*), ferritin light chain (*FTL*), and fibrinogen beta chain (*FGB*) expressed by hepatocytes; immunoglobulin kappa constant (IGKC) and heavy constant gamma (*IGHG1*) genes related to plasma cells, and *CD74* expressed by B cells (Figure 1C). Using established cell markers (Supplementary Data 2) in addition to unbiased clustering, we identified all major hepatic cell populations (Figure 1C) at higher resolution. As such, we were able to distinguish differentiated HSC (high *IGFBP7* and *COL1A1*) from their terminally activated forms, myofibroblasts (high *DCN, COL3A1, COL1A2*); separate sinusoidal endothelial cells (*PECAM1, STAB2, ADAMTS1*) and vascular endothelial cells (*ENG*, both populations VWF-positive). As expected, the strongest ALB signal was localized in the non-fibrotic parenchyma (Figure 1D). *PTPRC*, a pan-leukocyte marker, was mostly expressed on the fibrotic background. *VIM*, as a marker of mesenchymal lineage, was expressed throughout the tissue, with higher levels on the fibrotic background. VWF was found in the fibrotic areas, around major vessels, and in separate spots within hepatic parenchyma. Notably, VWF was previously described as a biomarker linked to endothelial dysfunction and portal hypertension, increasing in more advanced cirrhosis^12^.

The majority of the annotated cells in the marker-driven approach were identified as hepatocytes (∼64.1% of the total cell numbers), followed by NPC (∼21%, including cholangiocytes) and immune cells (∼15.1%; Figure 1E). Within the former two populations, fibroblasts and HSC comprised 75.3% of NPC, while the most abundant immune cell population consisted of plasma cells (33.3%). Application of a CellTypist model trained on human liver cells allowed us to increase cell annotation resolution, identifying numerous subclusters (Figure 1F, Supplementary Data 3). These subclusters included two NK cell types, non-inflammatory macrophages, hepatocytes, and endothelial cells, in regard to their zonation, various monocytes, different T cells, and dendritic cells (DC).

The NK types varied in *CD16* expression. Using the reference atlas of NK cells^13^, we found that our *CD16*^+^ NK cells subtype had features of mature NK cells with *TGFB1* and prostaglandin D2 synthase as the top upregulated genes. We suggest that cells of these clusters were involved in NK-endothelial interactions due to abundant expression of *ADAMTS1* and *ADAMTS4* (suppression of angiogenesis via eNOS-VEGF pathway^14^). In turn, CD16^-^ cells showed upregulation of *IRF8*, which was previously linked to clonal expansion^15^.

Non-inflammatory macrophages were distinguished by higher expression of the classical marker *MARCO*. Other significant differentially expressed genes (DEGs) included the anti-inflammatory cytokine *CD5L* and anti-inflammatory marker *TIMD4*^16^.

Hepatocytes annotated as periportal had high levels of portal markers *BMP7* and *HAL*^17,18^ and hemoglobin gene *HBZ*. Pericentral hepatocytes’ marker was *SLCO1B3*^19^. These cells overexpressed tuftelin 1 (*TUFT1*), previously linked to lipogenesis, focal adhesion and hepatocellular carcinoma development^20^. Those hepatocytes not belonging to either of these two groups had overexpression of dihydrofolate reductase (*DHFR*) and fibroblast growth factor receptor 2 (*FGFR2*), with the latter shown essential for liver regeneration in experimental models^21^.

STAB2^+^ pericentral LSEC had upregulation of *FGF23*, linked with endothelial dysfunction through the reduction of NO availability^22^. Conversely, periportal LSEC had upregulation of *CLEC14A* involved in cell adhesion^23^.

Tem/Temra T cells had prominent upregulation of cytotoxicity markers *GNLY, GZMH, NKG7*. Notably, they also had upregulation of *SLC16A7*, an indicator of active monocarboxylate metabolism^24^. This was different from Tem/Trm cytotoxic T cells with top upregulated markers *GZMK, THY1*, and *PRDM16*, a regulator of differentiation^25^. MAIT cells were positive for marker gene *SLC4A10* and expressed high levels of *IL7R* and *ITGA2*, which are involved in T cell-mediated inflammation and cell adhesion.

Further, two DC populations were identified, where DC1 had more pronounced expression of *CD74* and *TMSB4X*, and DC2 was negative for *IRF8*.

These results demonstrate that HD spatial transcriptomics allows for marker-based profiling of known cell types in a diseased liver and high-resolution clustering due to application of pre-trained machine learning models. Next, we evaluated the spatial expression of cell signatures related to immune infiltration and identified their diverse localization (Figure 1G). For example, pan-T cell signature (*CD3D, CD3E, CD3G, CD2*) was mainly abundant mainly in the fibrotic regions. In turn, the pan-macrophage signature, comprising cell and pro-inflammatory markers (*CD68, CD163, IL1B, IL6, MMP7, MMP9, IL10*), was widespread in both parenchyma and fibrotic areas – in contrast to the tissue-resident macrophage marker *MARCO* linked with non-inflammatory macrophages, not detected in fibrotic niches (Supplementary Figure 1B). The DC signatures originated from DEGs of the two subtypes identified as discussed above. Signature genes of DC1 were CD1D, *CLEC10A, LGALS2, SAMHD1, H3F3A*, whereas DC2 had upregulation of *TIMP3 CD1C, XCR1, CLEC9A, IDO1, WDFY4* (Figure 1H). Both signatures co-localized with fibrosis, and DC2 was more prominently expressed.

In summary, analysis of spatial features in ALD at ultra-high resolution (2µm bins) allowed the identification of major cell populations and their subtypes. It also allowed us to demonstrate the diversity of immune infiltrating cells in ALD, showing the presence of the pro-inflammatory macrophages in all tissue, with limited involvement of anti-inflammatory *MARCO*+ macrophages, which were not detected in the fibrotic areas, and differences in DC signatures.

### In-depth analysis of tissue niches identified *WNT4, RCAN3*, and other markers linked to fibrosis

To investigate zonation-related differences in cell abundances and expression patterns, we annotated the sample according to three types of regions: fibrotic (presence of the septae), vascular (portal triad or central vein), and parenchymal (neither of the two) niches. First, we confirmed the presence of the major cell populations in each niche (Figure 2A). As was defined from these areas, the highest cellular diversity was observed on the background of fibrotic niches and around major vessels.

**Figure 2.**
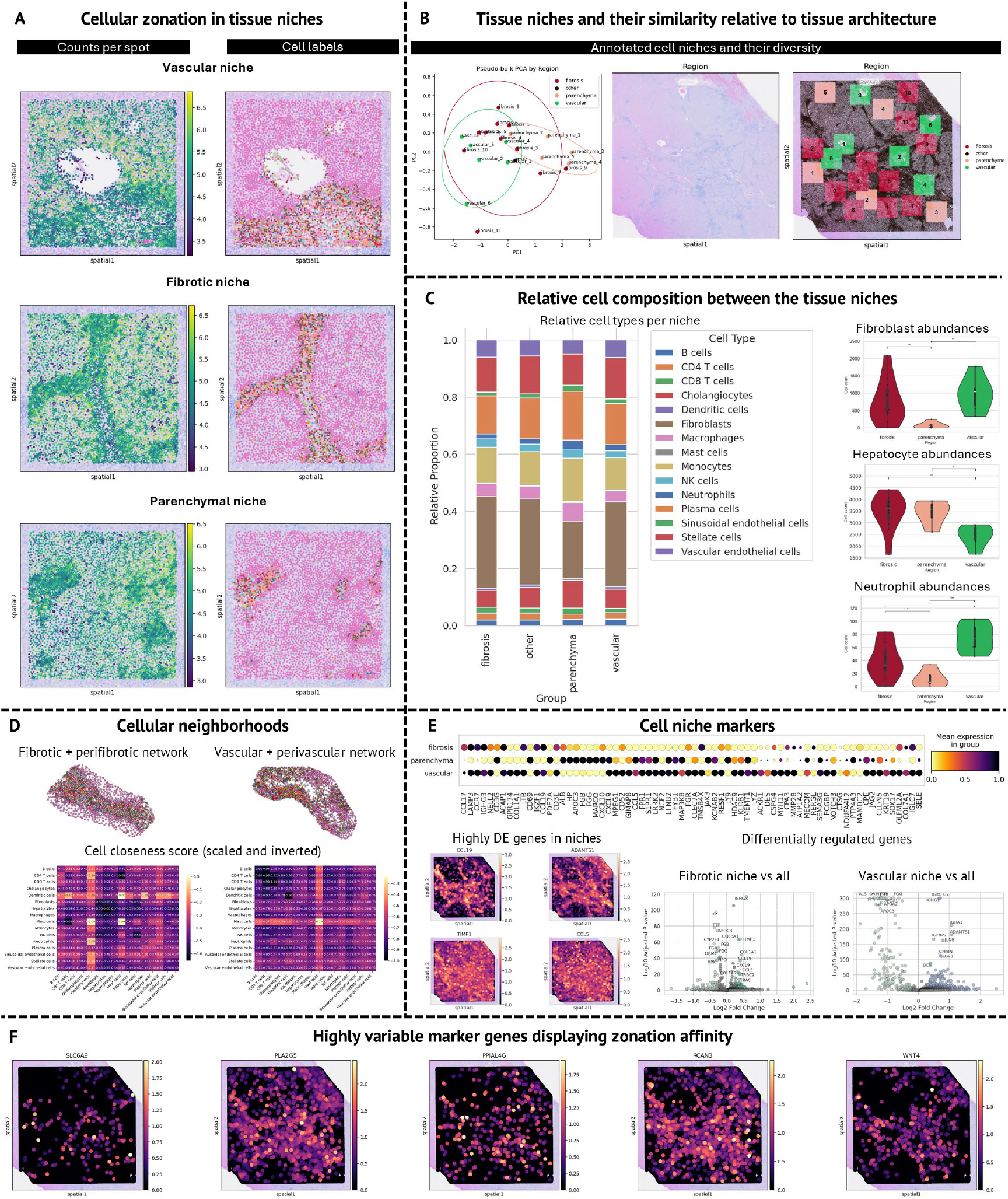
Tissue niches displayed various transcriptomic features in alcohol-related liver disease. **(A)** Representative annotation of vascular, fibrotic and parenchymal niche, color-coded with cell identities. **(B)** Principal component analysis of areas’ pseudo-bulk transcriptomes showing grouping of respective zones. An H&E stained and annotated slide is displayed for representation of the annotated liver regions. **(C)** Relative numbers of the identified cell populations between different zones (excluding hepatocytes). See also Supplementary Data 4. Selected cell populations are displayed to demonstrate significant differences between various cell niches. **(D)** Cellular neighborhood networks for fibrotic and vascular niches. Representative networks are visualized. The heatmaps indicate the closeness score, which we define as negative scaled average distance between two cell types. Brighter colors indicate that the cells are more often found in close contact between a niche and its surrounding space. **(E)** A dot plot displaying significantly differentially expressed genes between the ALD niches defined above, on pseudo-bulk level. Top differentially expressed genes are presented, together with volcano plots showing results of the differential testing of the fibrotic and vascular niches. **(F)** Marker genes which are, by combined spatial autocorrelation index, are selected as the markers with the highest affinity for fibrotic and vascular niches.

Next, we grouped all cellular transcriptomes at the pseudo-bulk level to investigate the baseline expression differences between the defined regions (Figure 2B). Principal component analysis showed a clear separation of parenchymal and vascular niches, while fibrotic niches had transcriptomic features similar to both of them, with a tendency to group better with vascular zones.

Across the three liver niches, cell type distributions exhibited marked heterogeneity (Supplementary Data 4). ANOVA analyses revealed significant differences for several populations, including cholangiocytes (p=0.020), fibroblasts (p=0.027), hepatocytes (p=0.016), macrophages (p=0.015), monocytes (p=0.008), NK cells (p=0.042), neutrophils (p<0.001), plasma cells (p=0.0136), sinusoidal endothelial cells (p=0.004), stellate cells (p=0.005), and vascular endothelial cells (p=0.015). Notably, parenchymal regions consistently harbored lower cell counts compared to fibrotic and vascular compartments. For example, fibroblasts were highly enriched in fibrotic areas (8,287 cells total) versus parenchyma (407 cells; fibrosis vs. parenchyma p=0.00485) (Figure 2C). Differentiated HSC showed similar pattern: enriched in fibrotic and vascular niches as compared to parenchyma (3,161 total in fibrotic niches, 2,865 in vascular; both p<0.05 versus parenchyma with 220 cells). Monocyte numbers were substantially higher in fibrosis (3,242 cells) and vascular regions (2,227 cells) than in parenchyma (307 cells; p<0.001 and p=0.008, respectively). Neutrophils were also significantly elevated in fibrotic (439 cells) and vascular (450 cells) niches relative to parenchyma (64 cells; fibrosis vs. parenchyma p=0.016, vascular vs. parenchyma p<0.001), and plasma cells followed a similar trend (Figure 2C). T cell subsets, although not significant by overall ANOVA (CD4 p=0.113; CD8 p=0.158), were reduced in parenchyma relative to fibrosis (CD4: p=0.025; CD8: p=0.027) and vascular areas (CD4: p=0.017; CD8: p=0.017). Dendritic cells showed a marginal overall difference (p=0.054) but were significantly lower in parenchyma than in both fibrosis (p=0.022) and vascular regions (p=0.008). Similarly, B cells were depleted in parenchyma (43 cells) compared to fibrosis (486 cells; p=0.0323) and vascular compartments (424 cells; p=0.007). Collectively, these findings underscore distinct cellular landscapes across liver niches - with fibrotic and vascular compartments generally enriched for immune and stromal populations relative to the non-fibrotic parenchyma. While investigating cellular abundances, we evaluated cellular neighborhoods to understand whether cellular proximity was varied between fibrotic and vascular niches. We found that the inflammatory infiltrate was more homogeneously distributed on the fibrotic septae, whereas vascular neighborhoods contained infiltration-free regions (Figure 2D). Moreover, we noted that mast cells, DCs, and NK cells were the closest neighbors in the perifibrotic niches, while only mast cells were frequent neighbors in the perivascular niche, followed by CD8 T cells with a more prominent proximity score difference (Figure 2D), suggesting the involvement of these cell identities in fibrotic processes.

Cellular markers differentially expressed between the liver zones were identified to further characterize the cellular composition within the three predetermined niches (Figure 2E, Supplementary Data 5). On the pseudo-bulk level, we established that a *CCL19, ACAP1, IGHG3, LAMP3*, and *RERGL* signature belonged to the fibrotic niche (Figure 2E). The vascular niche was strongly linked to *CXCL9, MPEG1, CD52, GIMAP8*, CLEC7A, and *NCF2* expression. As expected, parenchyma non-affected by fibrosis contained mostly hepatocyte-related expression as its signature prioritized expression of *ALB, HP, APOC3, FGB*, and others. Within the fibrotic environment, genes involved in matrix remodeling (*TIMP1, COL1A1, COL3A1*) and immune activation (*IGHG3, CCL19, CXCL9, CCL5*, various T cell receptor genes) were the most prominently differentially upregulated (Figure 2E). *ADAMTS1, DCN* and genes related to immunoglobulin chains were the most upregulated in the vascular areas (Figure 2E).

Notably, we found that *EGFL7*, an inhibitor of NF-kB and macrophage-endothelial adhesion^26^, was only upregulated in *MARCO*^+^ macrophages in parenchymal zones (Supplementary Data 6). Moreover, the *CCL19, CXCL9*, and *EGR1* subset was enriched in T cells, B cells, and DC populations in the fibrotic zones (Supplementary Figure 2B). *IFI27*, a member of *RCAN1*/*VEGFA* pathway linked to angiogenesis^27^, was upregulated in LSEC of parenchymal regions but not in those detected in other zones (Supplementary Figure 2C). These examples indicate that cell populations expressing identical sets of marker genes display different cellular transcriptomes based on the location within the tissue, thereby potentially mediating different effects dependent on the tissue environment they localize within. Following these findings, we calculated the spatial properties of the highly variable genes in our obtained dataset. We identified a subset of markers highly linked to the fibrotic regions, such as *WNT4, RCAN3*, and others (Figure 2E, Supplementary Data 7). The functional relevance of these markers will be further validated in the following paragraphs.

### The spatial interactome comprised CCL19-CCR7, DLL4-NOTCH3, and other signaling pathways in fibrotic and vascular niches

Following our findings on niche-specific markers and expression patterns, we investigated whether cellular residence in fibrotic or vascular niche itself drives interactome differences in ALD fibrosis. We noted that combined incoming and outgoing interactions, obtained by analysis of lig- and-receptor expression patterns, were lower in the parenchymal niche compared to fibrosis, and the highest scores were observed in the vascular niche (Figure 3A). LSEC, mast cells, and DC of the vascular niche had the highest interaction scores, while in the fibrotic microenvironment, CD4 T cells and DC displayed the highest number of interactions (Figure 3A).

**Figure 3.**
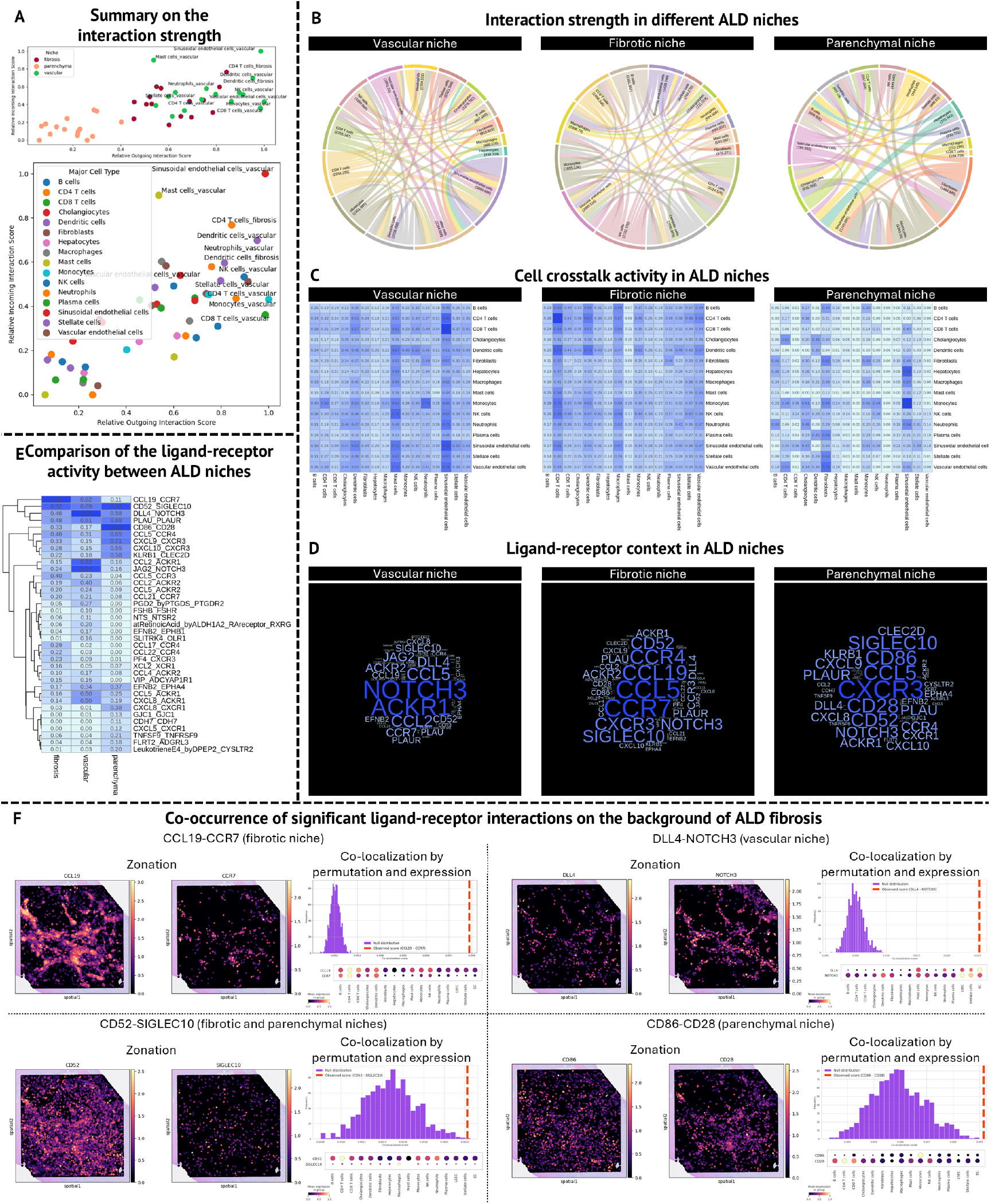
Hepatic ligand-receptor interactome during ALD in the spatial context. **(A)** Summary on the interaction strength between ALD niches and on the scale of separate cell types. The labels are displayed for the upper tertile. **(B)** Interaction maps showing differences in the incoming and outgoing interactions (as a combined score) between three ALD niches. **(C)** Activity of the ongoing cellular crosstalk, as identified by ligand-receptor interactions, in three ALD niches. **(D)** Ligand-receptor context, displaying the most highly scored ligands and receptors for each ALD niche. **(E)** Comparison of the ligand-receptor activity between the ALD niches. **(F)** Co-occurrence of significant ligand-receptor interactions on the background of ALD fibrosis. Expression of the most active ligand and receptor pairs are shown, together with their co-localization score and cellular attribution.

The vascular niche had the highest overall interaction strength across significant ligand-receptor pairs (sum of CellPhoneDB interaction scores 27789), with ∼35% more interactions than in the fibrotic area. Both niches had more than twice the number of interactions than occurring in the non-fibrotic parenchyma (CellPhoneDB score 10388; Figure 3B). The leading ligand-receptor activity in the vascular niche included crosstalk between LSEC, vascular endothelial cells, CD8, and CD4 T cells (Figure 3C, Supplementary Data 8). The fibrotic niche, in turn, had leading interactions between CD4 and CD8 T cells, DC, monocytes, and NK cells. This was distinct from the parenchymal landscape, where the highest crosstalk occurrence happened between monocytes, LSEC, hepatocytes and vascular endothelial cells.

We observed distinct ligand-receptor contexts in these microenvironmental contexts: the vascular niche had a predominance of *NOTCH3, ACKR1, CCL5, CCL2, JAG2* (Figure 3D). In the fibrotic niches, chemokines *CCL5* and *CCL19* were ligands with the most potent activity, and *CCR7*/*CCR4* were such receptors. The prioritized parenchymal ligands and receptors were *CXCR3, CCL5, SIGLEC10*, and *CD86*. When evaluating interacting ligand-receptor pairs, we marked *CCL19*-*CCR7* and *CD52*-*SIGLEC10* (fibrosis and parenchymal), *DLL4*-*NOTCH3* (vascular), and *CD52*-*SIGLEC10* in the parenchymal environment accordingly (Figure 3E). The measured co-localization rate of all the prioritized interactions was above the null distribution for given ligands and receptors in all cases (Figure 3F).

Niche-specific interactome analysis in ALD revealed that the cellular microenvironment significantly shaped ligand-receptor crosstalk. The vascular niche displayed the strongest and most diverse interactions -primarily among endothelial cells and T cells - whereas fibrotic areas were dominated by immune cell communications, particularly involving CD4 T cells and DC. In contrast, non-affected parenchymal regions exhibited notably lower interaction scores, underscoring the distinct signaling landscapes across liver microenvironments.

### WNT4^+^ fibroblasts are key drivers of the CCL19 communication network in hepatic single-cell and single-nuclei validation

Next, we accessed single-cell and single-nuclei transcriptomic datasets, obtained from patients with ALD cirrhosis via transjugular liver biopsy^28^, to evaluate and validate our newly identified markers for ALD niches and their prioritized interactions. Integration into a single dataset was done, which identified all major cell types; as expected, hepatocytes were the most abundant cell population, followed by DC and fibroblasts (Figure 4A). The marker-driven cell annotation was consistent with our spatial transcriptomic dataset, identifying robust cellular signatures (Figure 4B). Importantly, we could validate the expression of all identified markers with high zonation patterns in our spatial analyses on the cells in this dataset (Figure 4C). Notably, some of these marker genes, including *PADI2, WNT4, B3GALT6*, and *TMEM240*, were expressed only by small (<20%) subpopulations of the respective cell types (Figure 4C). Thus, we further explored whether these zonation-defined markers can also serve as defining markers for functionally active cell populations.

**Figure 4.**
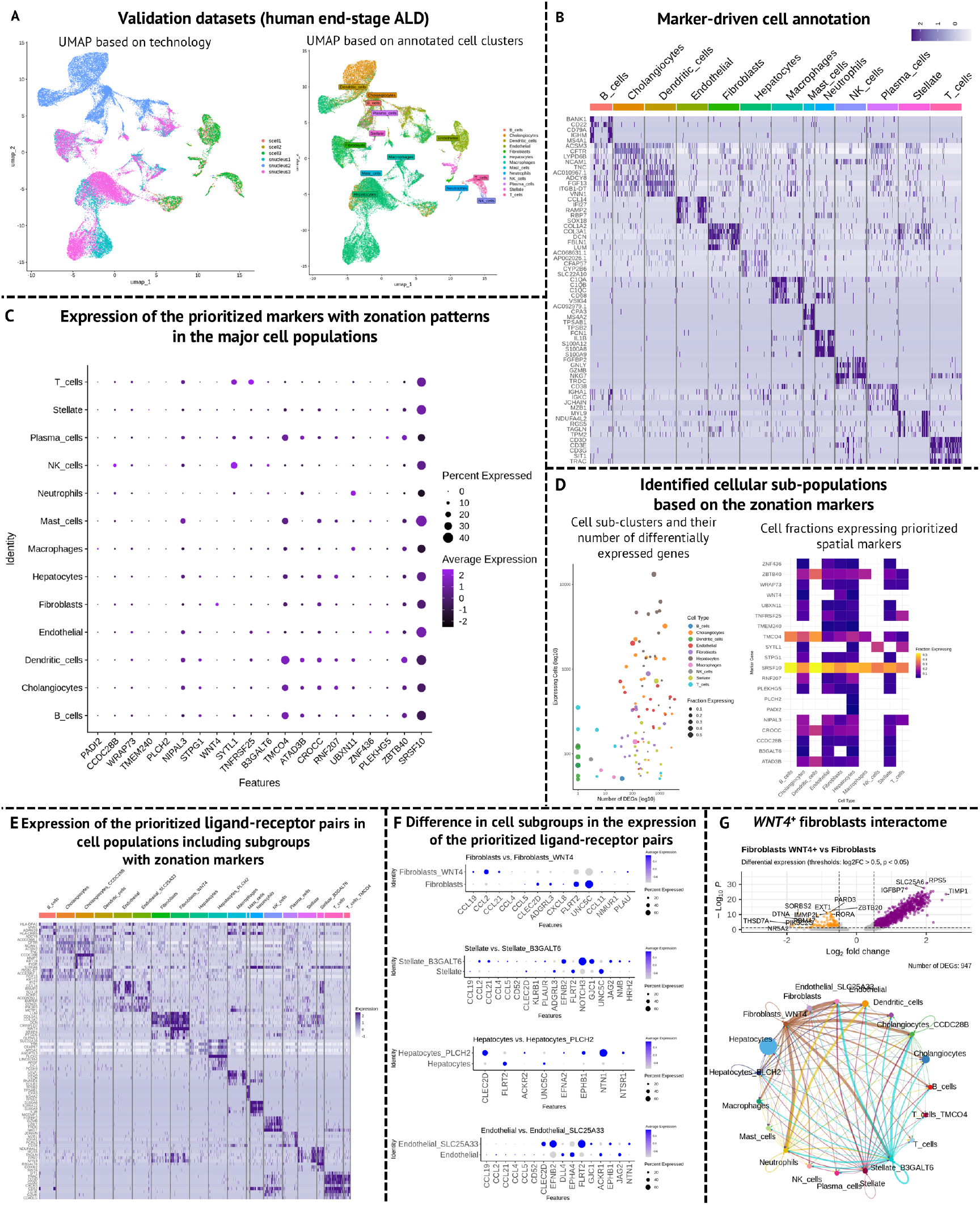
Validation of the zonation markers and spatial interaction on single-cell and single-nuclei datasets in ALD. **(A)** UMAP dimensionality reduction displaying datasets split by technology (left) and cell type annotation (right), post-integration. **(B)** Top five markers attributed to each of the cell clusters. **(C)** Expression of the highest prioritized zonation markers (see also Figure 2) within cell types of the validation dataset. **(D)** Ability of the cell markers to define functional cell subtypes, as defined by >1000 DEGs. **(E)** Heatmap showing expression of the prioritized ligand-receptor pairs within the major cell types, but also within the most transcriptomically active cell subtypes. **(F)** Comparison of the identified cell subtypes to their major cell cluster in regard to the differential expression of the prioritized ligands and receptors. **(G)** Differential testing results between WNT4+ fibroblasts and their major cell type, as well as WNT4+ fibroblasts’ contribution to *CCL19*-*CCR7* interactome in the validation dataset.

By selecting cells expressing one of the zonation markers from their respective significant cell populations, we found that, in most cases, these markers were important drivers of cellular heterogeneity (>1000 DEGs in cell sub-populations; Figure 4D). The hepatocytes cluster had the highest number of transcriptomically distinct subgroups, while the zonation markers did not drive DC diversity (Figure 4D). We identified robust populations of cholangiocytes, endothelial cells, fibroblasts, hepatocytes, stellate, and T cells based on the most specific spatial markers (*CCDC28B, SLC25A33, WNT4, PLCH2, B3GALT6, TMCO4* respectively; Figure 4E). Furthermore, we evaluated the expression of the prioritized interactions in the newly formed cell subpopulations. We found that *WNT4*^+^ fibroblasts, *B3GALT6*^+^ HSC, *PLCH2*^+^ hepatocytes, and *SLC25A33*^+^ endothelial cells varied in ligand-receptor expression from their respective major populations. *WNT4*^+^ fibroblasts had a signature of upregulated chemokines *CCL2, CCL4, CCL11, CCL19*, and *CCL21* (Figure 4F). *B3GALT6*^+^ HSC showed a remarkable upregulated on *NOTCH3, GJC1, EFNB2, JAG2*, and downregulation of *ADGRL3* and *FLRT2. PLCH2*^+^ hepatocytes had higher CLEC2D, NTN1, and EPHB1 levels and downregulation of *FLRT2* and *UNC5C*. Finally, *SLC25A33*^+^ endothelium had a signature of *CLEC2D, EFNB2* and *FLRT2* (up), *ACKR1*, and *EPHA4* (down).

We found that fibroblasts expressing *WNT4*, the marker highly linked to the fibrosis zones in our spatial dataset, had 947 DEGs versus their main cell cluster (Figure 4G). Their distinctive genes included *TIMP1* and *IGFBP7*, hinting at a more prominent role in extracellular matrix remodeling, and *SLC25A6*. We also observed that this particular cell population was the most active participant in the chemokine interactome from the prioritized ligand-receptor pairs (Figure 4G).

Integrating single-cell and single-nuclei transcriptomics in decompensated ALD cirrhosis validated spatially zonated markers across major liver cell types and uncovered distinct, functionally active subpopulations. In particular, marker-defined groups - such as *WNT4*+ fibroblasts - exhibited marked transcriptomic heterogeneity and prominent ligand-receptor interactions, underscoring their potential roles in extracellular matrix remodeling and fibrosis.

## Discussion

Our study represents the first ultra-high resolution spatial transcriptomics dataset to dissect the hepatic cellular architecture at 2µm resolution in ALD cirrhosis. The spatially resolved data underscored marked heterogeneity between fibrotic, vascular, and parenchymal zones. Fibrotic and vascular niches were enriched for immune and stromal populations relative to parenchyma, where immune infiltration was markedly reduced. Notably, the preferential localization of pan-T cell markers and pro-inflammatory macrophage signatures in fibrotic areas contrasts with the confinement of *MARCO*+ tissue-resident macrophages to parenchymal regions. These observations suggest that fibrosis not only reflects a quantitative increase in immune cells but also a qualitative shift in the resident versus infiltrative immune landscape.

Differential expression analyses reveal that identical cell types assume distinct transcriptomic identities depending on their microenvironment. For instance, the enriched expression of extracellular matrix (ECM) remodeling genes in myofibroblasts (e.g. *COL1A2*) within fibrotic regions contrasts with the more quiescent profile of HSC in non-fibrotic parenchyma. Similarly, our identification of zonation markers such as *WNT4, RCAN3*, and *PADI2* in fibrotic niches implies that localized signaling cues drive phenotypic specialization. The segregation of dendritic cell subpopulations - with DC2, characterized by upregulation of *TIMP3, CD1C*, and *XCR1*, being more prominent in fibrotic areas - further emphasizes the impact of tissue context on cellular function and immune activation.

Our interactome analysis highlighted that the vascular niche exhibits the highest overall ligand–receptor interaction scores, driven primarily by crosstalk among endothelial cells and T cells. In contrast, fibrotic zones show intensified interactions among CD4/CD8 T cells and DC, with a particularly notable role for chemokine pairs such as *CCL19*–*CCR7*. This spatial specificity in ligand–receptor engagement suggests that microenvironmental context not only modulates immune cell recruitment but also shapes downstream signaling pathways critical for fibrosis progression. Such insights into the molecular dialogue between stromal and immune cells may offer targeted avenues for therapeutic intervention.

*CCL19*-*CCR7*, zonated on the fibrotic background and the top-scored interaction in the fibrotic niche, mostly involved crosstalk between various T cell populations (Figure 3F). This axis was previously described as a potential therapeutic target in T cell and fibroblast interactome in various etiologies^29,30^, yet, leaving space for further exploration in liver fibrosis.

*CD52*, predominantly expressed by CD4 T cells, was predicted as a ligand for *SIGLEC10* (found in macrophages and DC) in both fibrotic and parenchymal microenvironments. It was previously described that sCD52-SIGLEC10 interaction is a part of the regulatory axis, preventing further TCR activation and T cell proliferation by reducing phosphorylation of kinases like Lck and Zap70^31^. Hence, this interaction can potentially contribute to immune homeostasis in the background of the ALD immune infiltration.

*DLL4*-*NOTCH3* (LSEC/vascular EC/HSC axis) is part of the WNT5B pathway previously linked to pro-inflammatory activation in lymphatic endothelium^32^, to which LSEC is traditionally attributed. Notably, therapeutic modulation of *NOTCH3* was previously suggested in pulmonary fibrosis, and its inhibiting has led to its attenuation, effectively reducing the amount of αSMA-positive myofibroblasts^33^.

Little is known about the role of *CD86*-*CD28* (monocytes-CD4 T cells axis) in liver fibrosis. Overall, this signaling was described as having an ambiguous role: depending on the *CD28*-expressing CD4 T cell subtype, it can either enhance Treg survival or promote T cell activation^34^. Moreover, CTLA-4 was described as a competitor for inhibitory CD28 co-stimulation (at higher affinity than CD86), hinting towards the need to further evaluate the role of therapeutic modulation of CD86 directly or via CTLA-4 in the context of ALD fibrosis^34^.

Integration with single-cell and single-nuclei datasets from decompensated ALD cirrhosis validates the spatially defined markers and uncovers functional subpopulations within major cell types. Of particular interest, *WNT4*+ fibroblasts emerge as key drivers of the chemokine interactome, exhibiting a transcriptomic signature enriched for *CCL2, CCL4, CCL11, CCL19*, and *CCL21*. The distinct differential expression in these fibroblasts - encompassing regulators of ECM remodeling such as *TIMP1* and *IGFBP7* - positions them as central mediators in fibrotic remodeling. This subpopulation’s heightened interaction with immune cells implicates *WNT4*+ fibroblasts in orchestrating the profibrotic microenvironment and emphasizes the therapeutic potential of modulating fibroblast-driven signaling networks.

Of note, while our dataset provides an opportunity to investigate markers solely based on the transcriptomic level, requiring other modalities (hepatic proteome, secretome) in further studies, integrating spatial profiling with single-cell and single-nuclei transcriptomics enabled us to delineate distinct immune, stromal, and parenchymal compartments within the cirrhotic liver, thereby advancing the understanding of niche-specific signaling and cellular interactions in fibrosis. With the newly identified ALD niches markers and key ligand-receptor interactions, we suggest targets for further investigation as biomarkers and in the context of therapeutic modulation. Further, it remains to be understood how our identified markers and cell populations vary during the natural course of the disease and whether they can be beneficial for monitoring fibrosis regression.

## Acknowledgements

We acknowledge the valuable contribution of the patients recruited to the LIVERMATRIX study to donate biospecimens. The transplant team of the Vienna General Hospital is acknowledged for providing access to the highly valuable sample. Kerstin Zinober and the technical staff of the Core Facilities of the Medical University of Vienna are acknowledged for the sample pre-processing.

## Author contributions

**Oleksandr Petrenko**: Methodology (equal); Formal analysis (lead); Writing – original draft (lead); Writing – review and editing (equal); Data curation (lead). **Thomas Sorz-Nechay**: Methodology (equal); Writing – review and editing (equal). **Martin Bilban**: Methodology (equal); Writing – review and editing (equal). **Sophia Derdak**: Methodology (equal); Writing – review and editing (equal). **Lukas Van Melkebeke**: Validation (equal); Writing – review and editing (equal). **Jef Verbeek**: Validation (equal); Writing-review and editing (equal). **Wenxin Huang**: Validation (equal); Writing review and editing (equal). **Schalk van der Merwe**: Validation (equal); Writing – review and editing (equal). **Benedikt S Hofer**: Methodology (equal); Writing – review and editing (equal). **Benedikt Simbrunner**: Methodology (equal); Writing – review and editing (equal). **Thomas Reiberger**: Methodology (equal); Writing – review and editing (equal). **Tim Hendrikx**: Conceptualization (lead); Funding acquisition (lead); Methodology (equal); Writing – review and editing (equal).

## Methods

### Human sample acquisition and processing

The liver specimen was obtained from a patient with end-stage liver disease related to ALD enrolled in the LIVERMATRIX study (Medical University of Vienna, ethical committee approval 2318/2019). During the liver transplantation surgery, a portion of the caudate lobe of the explanted liver was sampled and fixed at room temperature in 10% formalin over 24 hours. Then, the sample was dehydrated and infiltrated with paraffin with following picrosirius red and α-SMA staining in compliance with the local SOPs for histological samples processing (Supplementary Data 1, 2). The wholeslide images were then analyzed in HALO (V3.3.2541.184, Indica Labs), and the percentage of collagen-positive staining (% CPA) and α-SMA-positive area were quantified with the Area Quantification module.

### Spatial transcriptomics acquisition with 10x Visium HD

A liver tissue sample from a patient with ALD (study code LM_030) was processed using the 10x Genomics Visium HD Spatial Gene Expression platform. A 5 µm FFPE section was placed on a standard glass slide followed by H&E-staining and imaging on an Olympus VS200 slide scanner at 20x magnification according to the protocol provided by 10xGenomics (CG000684). The CytAssist instrument (10xGenomics) was used to facilitate the transfer of transcriptomic probes from the standard glass slide onto the Visium HD capture area followed by library preparation. CG000684). Visium Human Transcriptome Probe Set v2.0 hybridization, probe ligation, slide preparation, probe release, extension, library construction, and sequencing followed the Visium HD Spatial Gene Expression Reagent Kits User Guide (CG000685). Sequencing was performed on an Illumina NextSeq 2000 with paired-end reads (43 cycles Read 1, 10 cycles i7, 10 cycles i5, 50 cycles Read 2). Space Ranger v3.0.1 was used to map FASTQ files to the human reference (GRCh38-2020-A), detect the tissue section, align the sequencing data to the H&E microscopy image as well as the CytAssist image, and produce gene-barcode matrices for further downstream analysis.

### Cell identification and annotation

The sample analysis was performed in a Python environment (v3.12.0) in anndata format^35^ using scanpy (v1.10.0)^36^ to handle filtration and pre-processing steps. During the filtration step, genes expressed in <3 bins, as well as the spots with no identified gene transcripts, were discarded. The sample was pre-processed using the bin2cell workflow^37^. During it, 2 um bins were expanded and joined into cells using nuclei coordinates on the sample’s H&E image with StarDist (v0.7.3)^38^ segmentation. Those bins that did not confidentially merge during this step were identified with bin2cell spatial clustering as secondary labels and joined with respective cells based on their underlying expression. The merged cells were then additionally filtered by the following: expressed genes > 20, number of bins > 5, and genes expressed in < 3 cells were discarded. For cell type identification, cells were first subjected to unbiased clustering using Scanpy’s implementation of Leiden algorithm, where optimal resolution was set as 0.5. The primary cell type identification was achieved using Scanpy’s implementation of the modular score of known marker genes (Supplementary Data 2). Further sub-clustering was achieved using CellTypist (v1.6.3)^39^ with the combined liver cell model.

### Hepatic niche identification and cell quantification

Niche identification was done in QuPath (v.0.5.1) as 50×50 pixel zones. Vascular niches were defined based on the classical signs of portal triads, fibrotic zones were selected based on the underlying picrosirius red staining. Zones without such signs were identified as parenchyma. During pre-processing, the dataset was filtered using pandas to exclude unwanted observations (regions non-labeled as niches). After filtering, a complete grid was constructed by merging unique replicates with all available cell type categories, ensuring that each combination of region, replicate, and cell type was represented. Cell counts were then computed and aggregated into contingency tables, from which relative proportions were derived by normalizing the counts over group totals. For statistical testing purposes, one-way ANOVA tests were performed using SciPy (v1.15.2) implementation to detect overall differences among regions, and pairwise comparisons were carried out using SciPy’s Welch’s t-test (via SciPy’s ttest_ind() function with unequal variances) for each cell type.

For niche neighborhood analysis, polygon geometries were extracted from each annotation, with MultiPolygon features decomposed into individual polygons while line geometries were excluded. Each polygon was buffered by 20 pixels to generate a surrounding peripheral region to get perifibrotic and perivascular zones. Cell-type compositions were computed by grouping cells by their assigned spatial neighborhood and calculating the frequency of each cell type, while pairwise intercellular distances were derived using a Euclidean distance matrix to obtain mean and median distances for celltype pairs. For spatial network analysis, cells from specific zones were filtered and connected using a KDTree query with a distance threshold of 80.0 units; network edges were aggregated by cell-type pair, globally scaled by inverting the mean distances (normalized by the global minimum and maximum), and multiplied by –1 to yield negative weights that facilitate downstream comparisons across cellular niches.

### Identification of the zonation markers

Zonation analysis was performed using Squidpy (v1.6.1)^40^. In brief, we ranked all genes using three methods. Moran’s spatial autocorrelation and Geary’s spatial auto-correlation were used as suggested by the authors of the workflow. Those genes meeting the FDR < 0.05 threshold following the Benjamini-Hochberg procedure (i.e., displaying spatial patterns) were subjected to the following analysis. Additionally, genes were ranked according to the Sepal score^41^. All three scores were scaled between 0 and 1, and merged to obtain the combined zonation score used further to evaluate genes displaying spatial expression patterns.

### Interaction analysis

We utilized interaction analysis in two contexts: for identification of spatially-significant interactions and using a standard single-cell-compliant approach. For spatial interactions, Squidpy’s implementation of the CellChatDB ligand-receptor permutation^42^ was utilized with threshold = 0.01 and n = 1000 permutations. Those interactions with P^adjusted^ < 0.05 were taken for further evaluation. We used LIANA+ (v1.4.0)^43^ interaction ranking in the consensus mode for the single-cell-compliant approach, with a minimum expression proportion for the ligands and receptors, including subunits, of 0.1. For further evaluation, the results were ordered by LIANA’s specificity and magnitude ranks. For co-localization analysis of the prioritized ligand-receptor pairs, we performed a permutation test to assess ligand-receptor co-localization. First, the observed co-localization score was computed as the average fraction of neighboring receptor expression in lig- and-positive spots, where each spot’s receptor contribution was normalized by its neighbor count. To generate a null distribution, receptor labels were randomly permuted 1,000 times. In each permutation, the receptor values were shuffled, the weighted sum over neighbors was recalculated via matrix multiplication with the neighbor connectivity matrix, and the corresponding neighbor receptor fraction was determined for ligand-positive spots. The p-value was defined as the proportion of permuted scores greater than or equal to the observed score, while a Z-score was calculated by standardizing the observed score relative to the null distribution’s mean and standard deviation.

### External validation of the predicted interactions

The single-cell and single-nuclei datasets previously obtained at the University Hospital Leuven (ethical approval S64744) were used in the validation experiments^28^. The datasets (n = 6) were pre-processed in Seurat (v5.2.0)^44^, and integrated using RPCA. The cell identity was annotated using module score calculation with established major liver cell makers we previously used in the spatial dataset (Supplementary Data 2). Then, the expression of the prioritized spatially-resolved markers and interactions was evaluated, with further cell subclustering by such markers. CellChatDB (v2.1.0, database v5.0.0) was utilized for the interaction analysis.

## Data availability

The original data produced for the manuscript, as well as the code used to perform the analysis, are available from the corresponding author upon reasonable request. Both data and the code will be released in a relevant public repository upon the manuscript acceptance decision. Supplementary files are available via https://doi.org/10.5281/zenodo.15030351.

## Abbreviations

Abbreviation: Full name
ALD: Alcohol-related liver disease
ALT: Alanine transaminase
AST: Aspartate transaminase
ANOVA: Analysis of variance
ACAP1: ArfGAP with Coiled-Coil, Ankyrin Repeat and PH Domain 1
ADAMTS1: A disintegrin and metalloproteinase with thrombospondin motifs 1
ALB: Albumin
APOC3: Apolipoprotein C3
BMP7: Bone morphogenetic protein 7
B3GALT6: Beta-1,3-galactosyltransferase 6
CCDC28B: Coiled-coil domain containing 28B
CCL2: C-C motif chemokine Ligand 2
CCL4: C-C motif chemokine Ligand 4
CCL5: C-C motif chemokine Ligand 5
CCL11: C-C motif chemokine Ligand 11
CCL19: C-C motif chemokine Ligand 19
CCL21: C-C motif chemokine Ligand 21
CD1C: Cluster of differentiation 1c
CD1D: Cluster of differentiation 1d
CD2: Cluster of differentiation 2
CD3D/E/G: CD3 (T-cell receptor complex subunits: delta/epsilon/gamma)
CD16: Cluster of differentiation 16 (FcγRIII)
CD28: Cluster of differentiation 28
CD52: Cluster of differentiation 52
CD68: Cluster of differentiation 68
CD74: Cluster of differentiation 74
CD86: Cluster of differentiation 86
CLEC10A: C-type lectin domain family 10 member A
CLEC14A: C-type lectin domain family 14 member A
CLEC2D: C-type lectin domain family 2 member D
CLEC7A: C-type lectin domain family 7 member A
CLEC9A: C-type lectin domain family 9 member A
COL1A1: Collagen type I alpha 1 chain
COL1A2: Collagen type I alpha 2 chain
COL3A1: Collagen type III alpha 1 chain
CTLA-4: Cytotoxic T-lymphocyte-associated protein 4
CXCL9: C-X-C motif chemokine ligand 9
CXCR3: C-X-C motif chemokine receptor 3
DC: Dendritic cells
DCN: Decorin
DHFR: Dihydrofolate reductase
DLL4: Delta-like canonical Notch ligand 4
EC: Endothelial cells
EGR1: Early growth response 1
EFNB2: Ephrin-B2
ENG: Endoglin
EPHB1: Eph receptor B1
EPHA4: Ephrin type-A receptor 4
FGB: Fibrinogen beta chain
FGF23: Fibroblast growth factor 23
FGFR2: Fibroblast growth factor receptor 2
FIB4: Fibrosis-4 score
FTL: Ferritin light chain
GIMAP8: GTPase, IMAP family member 8
GJC1: Gap junction protein gamma 1
GNLY: Granulysin
GZMH: Granzyme H
GZMK: Granzyme K
HSC: Hepatic stellate cells
HAL: Histidine ammonia-lyase
HBZ: Hemoglobin subunit zeta
IGFBP7: Insulin-like growth factor binding protein 7
IGHG1: Immunoglobulin heavy constant gamma 1
IGHG3: Immunoglobulin heavy constant gamma 3
IGKC: Immunoglobulin kappa constant
IL1B: Interleukin 1 beta
IL6: Interleukin 6
IL7R: Interleukin 7 receptor
IL10: Interleukin 10
IFI27: Interferon alpha-inducible protein 27
IRF8: Interferon regulatory factor 8
ITGA2: Integrin subunit alpha 2
JAG2: Jagged canonical Notch ligand 2
LAMP3: Lysosome-associated membrane protein 3
Lck: Lymphocyte-specific protein tyrosine kinase
LSEC: Liver sinusoidal endothelial cells
MARCO: Macrophage receptor with collagenous structure
MAIT: Mucosal-associated invariant T cells
MMP7: Matrix metalloproteinase 7
MMP9: Matrix metalloproteinase 9
MPEG1: Macrophage expressed gene 1
NF-kB: Nuclear factor kappa-light-chain-enhancer of acti-
vated: B cells
NCF2: Neutrophil cytosolic factor 2
NK: Natural killer cells
NTN1: Netrin 1
NO: Nitric oxide
NKG7: Natural killer cell granule protein 7
PADI2: Peptidyl arginine deiminase 2
PCA: Principal component analysis
PECAM1: Platelet endothelial cell adhesion molecule 1
PTPRC: Protein tyrosine phosphatase receptor type C
RCAN1: Regulator of calcineurin 1
RCAN3: Regulator of calcineurin 3
RERGL: RERGL Ras-like GTPase
SAMHD1: SAM domain and HD domain 1
SIGLEC10: Sialic Acid binding Ig-Like lectin 10
SLCO1B3: Solute carrier organic anion transporter family member 1B3
SLC16A7: Solute carrier family 16 member 7
SLC25A33: Solute carrier family 25 member 33
SLC25A6: Solute carrier family 25 member 6
SLC4A10: Solute carrier family 4 member 10
STAB2: Stabilin-2
Tem: Effector memory T cells
Temra: Terminally differentiated effector memory T cells Re-expressing CD45RA
Trm: Tissue-resident memory T cells
TIMD4: T-cell immunoglobulin and mucin domain containing 4
TIMP1: Tissue inhibitor of metalloproteinases 1
TIMP3: Tissue inhibitor of metalloproteinases 3
TMSB4X: Thymosin beta-4, X-linked
TMCO4: Transmembrane and coiled-coil domains 4
TMEM240: Transmembrane protein 240
TCR: T Cell Receptor
Treg: Regulatory T Cells
TUFT1: Tuftelin 1
WDFY4: WD repeat and FYVE domain containing 4
WNT4: Wingless-type MMTV integration site family member 4
WNT5B: Wingless-type MMTV integration site family member 5B
XCR1: X-C motif chemokine receptor 1
Zap70: Zeta-chain-associated protein kinase 70

---

**Supplementary Figure 1.**
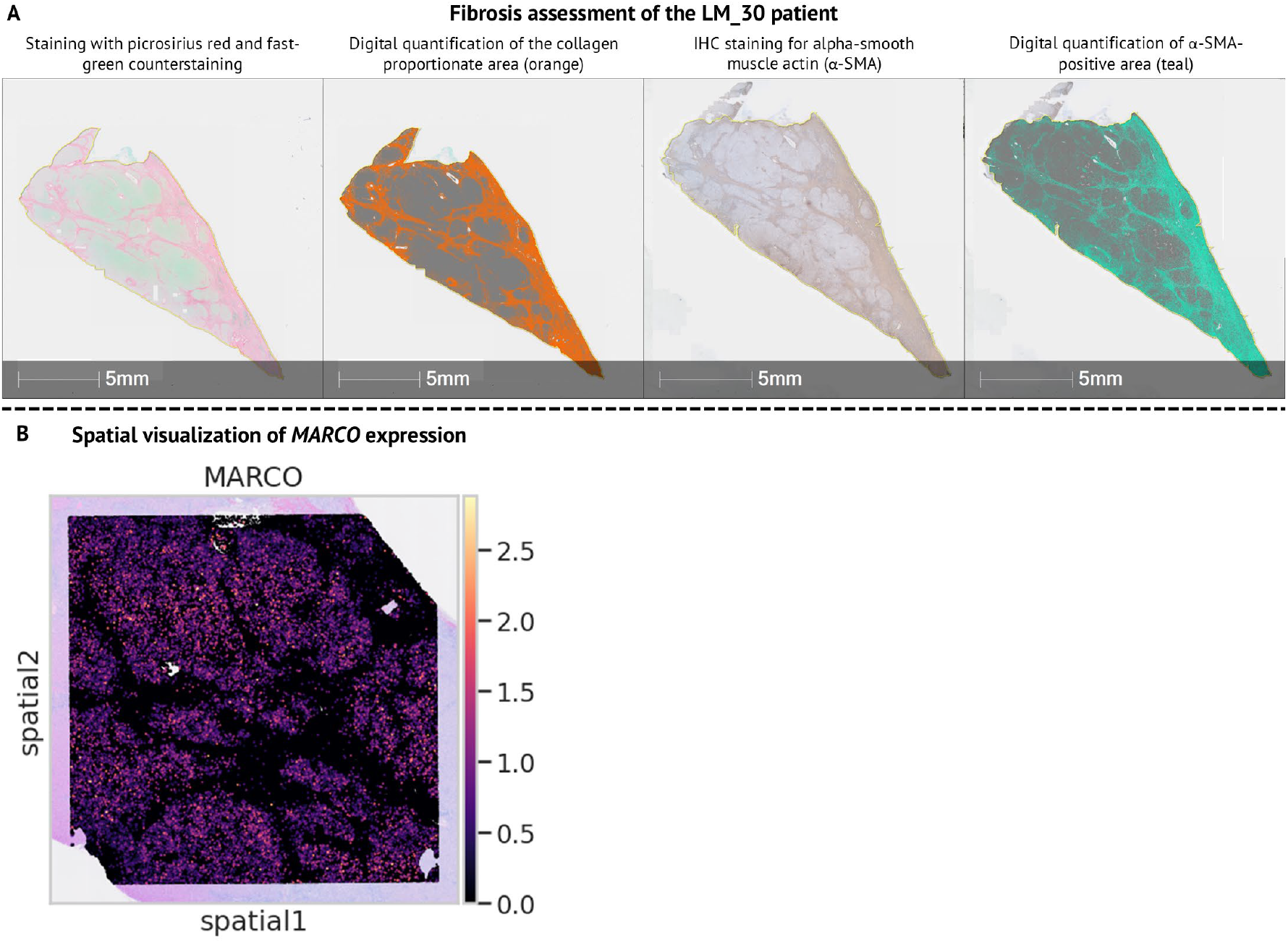
Fibrosis characterization of the included patient. **(A)** Staining with picrosirius red and for α-SMA. Digital quantification of the collagen abundance and α-SMA-positive hepatic stellate cells are displayed side by side. **(B)** *MARCO* expression pattern displaying presence of non-inflammatory macrophages outside of fibrotic septae.

